# Quantifying human adaptation to a novel split-belt walking condition after broad experience at different belt speeds

**DOI:** 10.1101/2025.08.25.671956

**Authors:** Zijie Jin, Jason Isa, Samuel A. Burden, Kimberly A. Ingraham

## Abstract

Humans can adapt their gait to minimize energy cost when given sufficient exposure to novel energy landscapes. However, it remains unclear whether broad experience in one energy landscape is sufficient to initiate continuous optimization when encountering similar but distinct conditions. In this study, we used visual biofeedback to guide 15 participants to broadly explore walking gaits with different step length asymmetries (SLAs) while walking on a split-belt treadmill with a large split-belt ratio. We subsequently tested how participants adapted their walking patterns during free exploration trials at the large split-belt ratio and at a new, smaller split-belt ratio. Our results showed that during guided exploration, participants were exposed to energetically favorable conditions. However, participants did not self-select walking patterns that minimized their metabolic cost during either free exploration trial. When first exposed to the smaller split-belt ratio, participants immediately adjusted their leg swing distances and belt contact times while maintaining similar step lengths. Throughout this trial, they continued adapting by significantly increasing step lengths on the fast belt. Taken together, our results suggest that participants actively modified their gait strategy when exposed to an energy landscape distinct from the one in which they gained broad experience.

## Introduction

Walking on a split-belt treadmill, in which one belt moves faster and one belt moves slower, is a task that has commonly been used to study human locomotor adaptation and learning ^1–5^. Walking on a split-belt treadmill perturbs the person’s usual, symmetrical gait and introduces errors, a process believed to initiate motor adaptation^4,6–9^. Recent studies have suggested that walking on a split-belt treadmill can theoretically reduce the energetic cost of walking below that of tied-belt walking^10–12^. Since it is metabolically more costly for muscles to do positive work than negative work^11,13,14^, humans can save energy if they reduce the amount of positive work done by their muscles. Such a reduction can be achieved by the split-belt treadmill doing positive work on the human. This occurs when humans adopt a particular gait pattern—that is, walk with a positive *step-length asymmetry* (SLA). SLA is defined as the normalized difference in step length between consecutive steps taken on the faster belt and those on the slower belt. Thus, if humans take a larger step forward on the fast belt relative to the slow belt, the work performed by the treadmill on the person can reduce the positive work required by the person’s muscles and ultimately reduce their metabolic cost^11^. However, this energetic benefit is not guaranteed when walking with a positive SLA, because humans may choose to dissipate this energy by performing additional negative work. Therefore, humans must learn to reduce the positive work done by their muscles to reduce their metabolic cost.

It is well-understood that humans prefer to walk with gaits that minimize energy cost^15–17^. Accordingly, it is reasonable to expect that humans would be able to adapt their gait on a split-belt treadmill to adopt the energetically optimal strategy of walking with a positive SLA and accepting the positive work done by the treadmill^11,18^. *In silico* simulations have indeed demonstrated that machine-learning-powered agents (such as a reinforcement learning agent^19^ and a muscle-effort-minimizing agent^20^) will spontaneously adopt this gait pattern during a split-belt walking task. However, such observations were not consistent across *in vivo* studies. While most experimental studies reported a gradual increase in the human’s SLA from initially negative to near symmetrical at steady state, the steady-state SLAs observed in many of them were not significantly more positive than the tied-belt baseline^3,21–25^. While a symmetrical gait incurs lower metabolic penalty in split-belt walking compared to gaits with negative SLA^11,21,22^, more energetic savings could be achieved by adopting a positive SLA.

One explanation for this phenomenon is that humans prefer walking with a symmetric gait, which typically minimizes energy cost during normal walking^26,27^. Prior research suggests that humans tend to stick to their existing preferred gait patterns, even in novel energy landscapes where adopting a different gait pattern could lead to higher energetic savings. Humans’ natural gait variability is not sufficient to initiate adaptation, and instead, humans only spontaneously optimize their gait patterns to minimize energy cost if they have some explicit, broad experience within the new landscape^28,29^. This has been observed for a variety of locomotion tasks with novel energetic landscapes, such as walking with a resistive exoskeleton^28,29^, an assistive mechatronic device^30^, and at varying inclinations^31^.

For split-belt treadmill walking, experiments indicate that humans require either enforced broad experience within the new energy landscape^11^, or very long exposure times (> 30 minutes)^18^, to learn that walking with positive SLA and accepting treadmill work can lower their metabolic cost. Once they have learned this strategy, however, they can quickly and spontaneously converge to energetically efficient gaits. Among these explanations, broad experience within the cost landscape is likely a key contributor to the process, as multiple studies have reported an absence of spontaneous adaptation to positive SLA despite overall protocol lengths >45 minutes^21,22^.

As previously stated, after sufficient exposure to a novel energy landscape, humans can quickly and spontaneously converge to energetically efficient gaits that differ from their typical symmetrical gait patterns. However, we do not yet know if broad experience within one novel energy landscape is sufficient to initiate continuous optimization in a new energy landscape with similar properties to the one they experienced. On one hand, it is possible that broad experience leads humans to developing a novel—yet fixed—gait strategy that is optimal in current landscape. Alternatively, broad experience could enable humans to not only adapt within the current landscape but also extrapolate beyond it and continuously adapt in response to perturbations to the landscape.

In this study, we ask whether and how people adapt the strategies that they previously developed through broad experience with one energy landscape when exposed to a similar yet distinct energy landscape. One way to create distinct energy cost landscapes within the split-belt treadmill walking paradigm is to alter the individual belt speeds while maintaining the same average speed across belts^22,24^. This can be quantified by the *split-belt ratio* (SBR), defined as the ratio of the fast belt speed to the slow belt speed^32^.

Some previous studies have investigated how humans adapt to split-belt walking at different split-belt ratios, but to our knowledge, these studies have not evaluated how participants adapt to a new split-belt ratio after they broadly experience walking at a different split-belt ratio. For example, Leech et al. compared how participants’ SLA changed as a result of different split-belt ratios, where the slow belt speed was kept constant and the fast belt speed was changed^24^. While this study involved multiple split-belt ratios, it focused on studying how people spontaneously adapt and re-adapt, rather than their response to guided experience. Stenum and Choi mapped the metabolic cost landscapes created by different split-belt ratios (maintaining the same average belt speeds) by constraining different parameters of participants’ gait^22^. This study focused on mapping out the participants’ energy cost landscapes at various gaits instead of analyzing participants’ self-selected response to a broad range of experiences.

In this study, we guided 15 participants walking on a split-belt treadmill to broadly explore an energy landscape defined by a large split-belt ratio and subsequently tested how they responded to walking at a new, smaller split-belt ratio with the same average belt speeds (Figure 1) . We adapted a previously described split-belt protocol with biofeedback^11,33^ for our study. During the *guided exploration* phase, we guided participants walking on the treadmill with a split-belt ratio of 3 to achieve seven distinct SLAs from −0.15 to +0.15 in increments of 0.05. Participants walked at each SLA for 6 minutes each. During the guided exploration phase, we displayed participants’ real-time foot position and encouraged them to match the target step lengths displayed on a tablet in front of them to achieve the desired SLA. We then removed the visual feedback and asked the participants to freely walk with a self-selected SLA in a *free exploration* trial at the same split-belt ratio of 3. Finally, participants completed a second free exploration trial at a different, smaller split-belt ratio of 2. During the walking trials, we measured participants’ spatiotemporal gait parameters and metabolic cost. We computed the work performed by the human legs and the treadmill using previously published methods^11,12,16,34,12,16^. To determine how participants adapt their gait when walking at a smaller split-belt ratio after guided experience on a larger split-belt ratio, we compared participants’ energetics and biomechanics between the two free exploration trials. Our novel contribution is the inclusion of the second free exploration trial at a smaller split-belt ratio than the guided exploration trials. This is to quantify whether, and if so, how participants adapted their gait to this new condition that they had not previously experienced.

**Figure 1.**
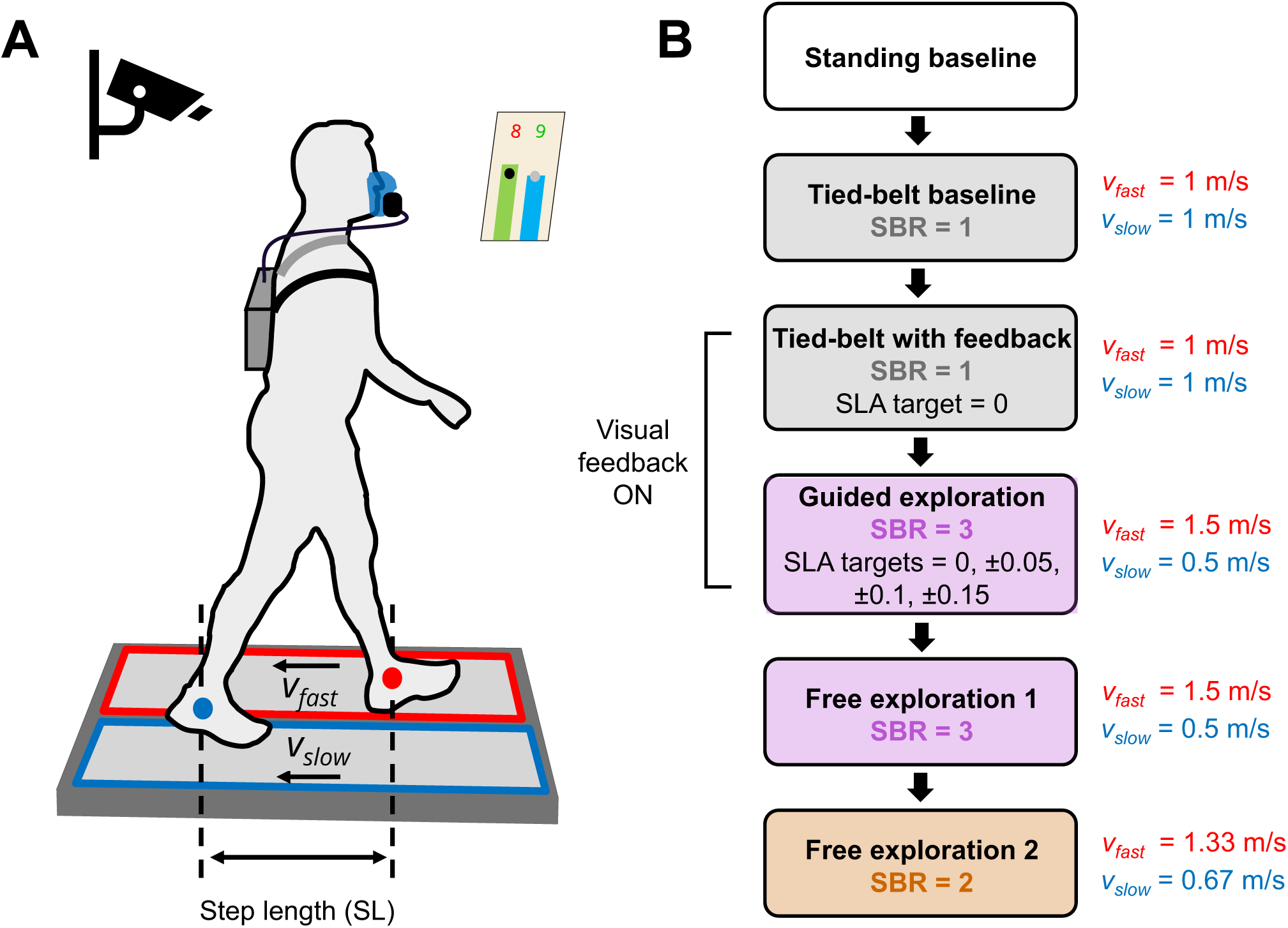
Overview of experimental protocol. (A) Fifteen participants walked on a split-belt treadmill with the left belt moving faster than the right belt. We placed motion capture markers on both lateral malleoli to determine the real-time position of each foot. (B) Participants completed eleven 6-minute walking trials. We first measured the participants’ baseline gait in a tied-belt trial, then added visual feedback to a second tied-belt trial (guiding participants to their baseline step length asymmetry, SLA) to familiarize the participants with the GUI. We next guided participants to walk with SLAs of −0.15, −0.1, −0.05, 0, 0.05, 0.1 and 0.15 using the GUI, at a split-belt ratio (SBR) of 3. Finally, we removed the visual feedback and had participants walk freely during two free exploration trials at SBRs of 3 and 2, respectively.

## Results

During the *guided exploration* trials, participants walked on the treadmill at a split-belt ratio of 3, guided by the visual feedback to walk at seven distinct step-length asymmetries (SLAs). We analyzed the work rate done by the human legs and the treadmill as well as the human metabolic cost as a function of SLA using linear mixed effect models (LMEM). The net work rate done by the human legs decreased linearly with SLA (−0.027 [−0.030, −0.025], p < 0.001) (Figure 2 A), and at positive SLAs, the human legs did net negative work (Figure 2 A). Conversely, the net work rate done by the treadmill on the human legs increased linearly with SLA (slope [95%CI] = 0.028 [0.025, 0.030], p < 0.001), and at positive SLAs, the treadmill did net positive work on the human (Figure 2 B). Human leg positive work rate decreased linearly with treadmill total work rate (slope [95%CI] = −0.546 [−0.671, −0.420], p < 0.001). (Figure 2 C). Participants’ metabolic cost linearly decreased with SLA (slope [95%CI] = −0.061 [−0.092, −0.031], p < 0.001) (Figure 2 D).

**Figure 2.**
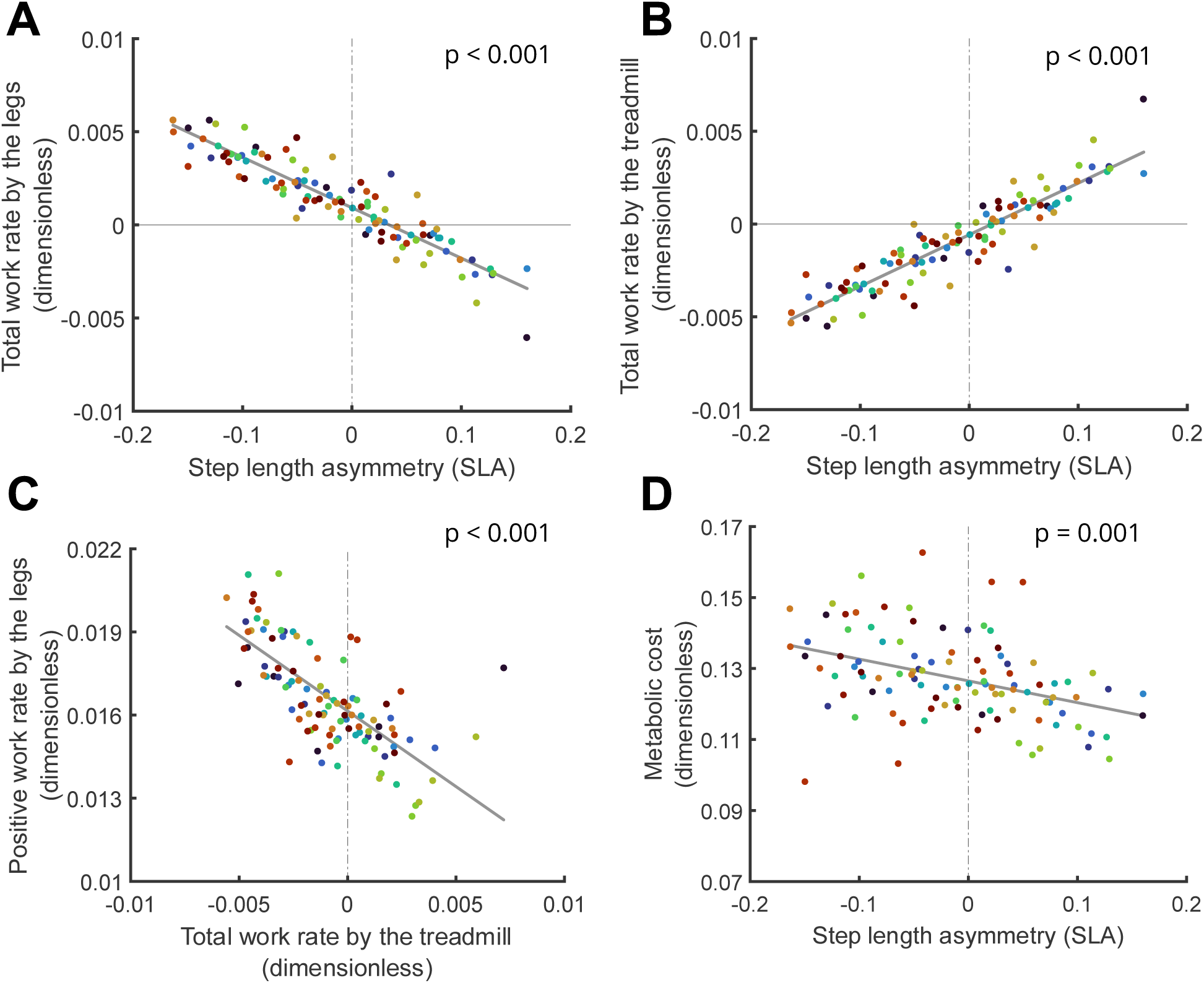
Linear mixed effect models of participants’ energetics vs. step length asymmetry (SLA) during guided exploration trials. Each data point represents a single guided exploration trial of a single participant, and different colors represent different participants. P-values indicate significance of the fitted slope. As SLA went from negative to positive, (A) participants’ total work rate done by legs decreased while, (B) total work rate done by treadmill increased. (C) The decrease in participants’ total leg work rate was linearly correlated with the increase in the treadmill’s total work rate. (D) Participants’ metabolic energy cost decreased as SLA increased from negative to positive.

During the *free exploration* trials, participants did not self-select SLAs more positive than their tied-belt baseline for either split-belt ratio, despite potential energy savings (Figure 3). We used paired student’s t-tests to compare self-selected SLAs between conditions. Average self-selected SLA across participants during the first free exploration trial (SBR = 3) was 0.010 ± 0.059 (mean ± SD). This SLA was not significantly different from participants’ baseline SLA on the tied-belt treadmill, which was −0.006 ± 0.026 (p = 0.37). Average self-selected SLA across participants during the second free exploration trial (SBR = 2) was 0.013 ± 0.052, which was not significantly different from tied-belt baseline (p = 0.22) or the first free exploration trial (p = 0.67) (Figure 3). We observed significant differences in metabolic cost between tied-belt baseline and two free exploration trials (p = 0.003, ANOVA) (Figure 4).

**Figure 3.**
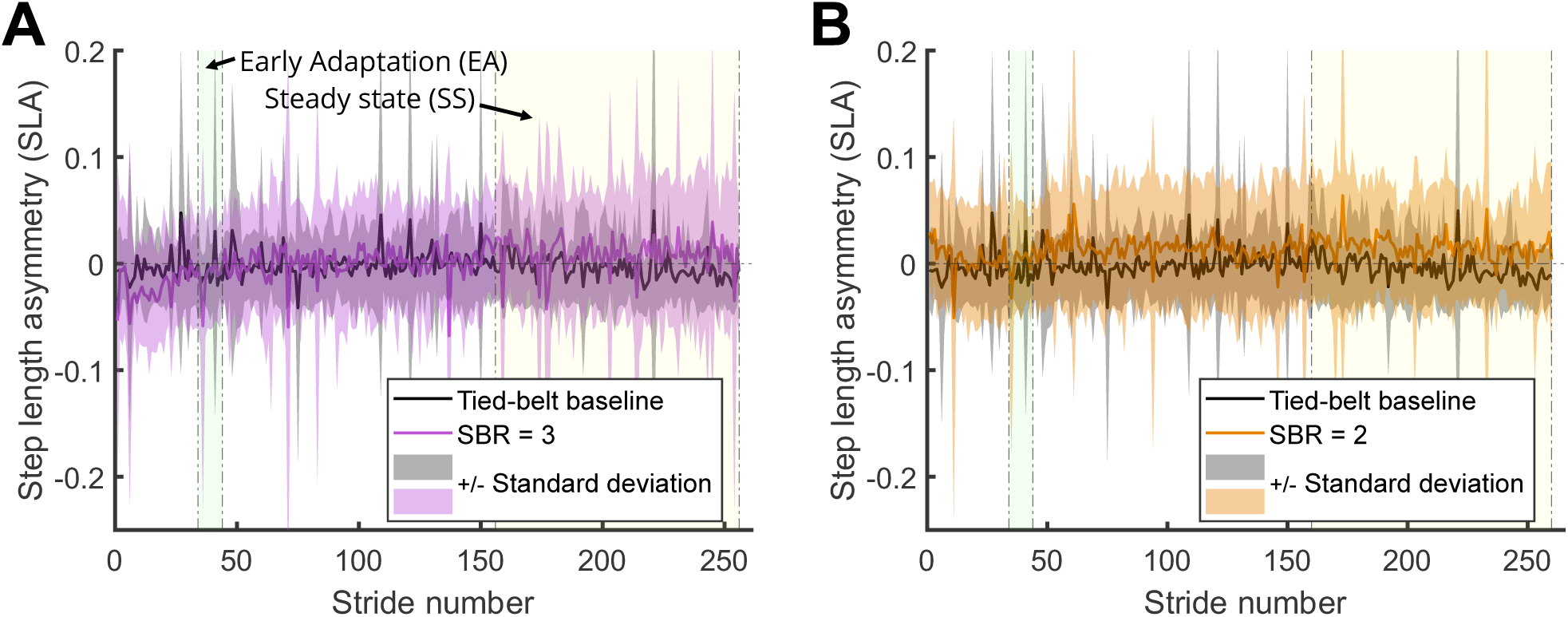
Stride-by-stride SLA comparison between tied-belt baseline and the free exploration trials with split-belt ratio of 3 (A) and 2 (B), averaged across participants. Purple, orange, and grey shaded areas indicate standard deviation of each stride’s SLA across participants. The green shaded area indicates the“early adaptation” period (strides 34 – 44), and the yellow shaded area indicates the “steady-state” period (last 100 strides of each trial). Neither free exploration trial’s steady-state SLA was significantly higher than that of tied-belt baseline.

**Figure 4.**
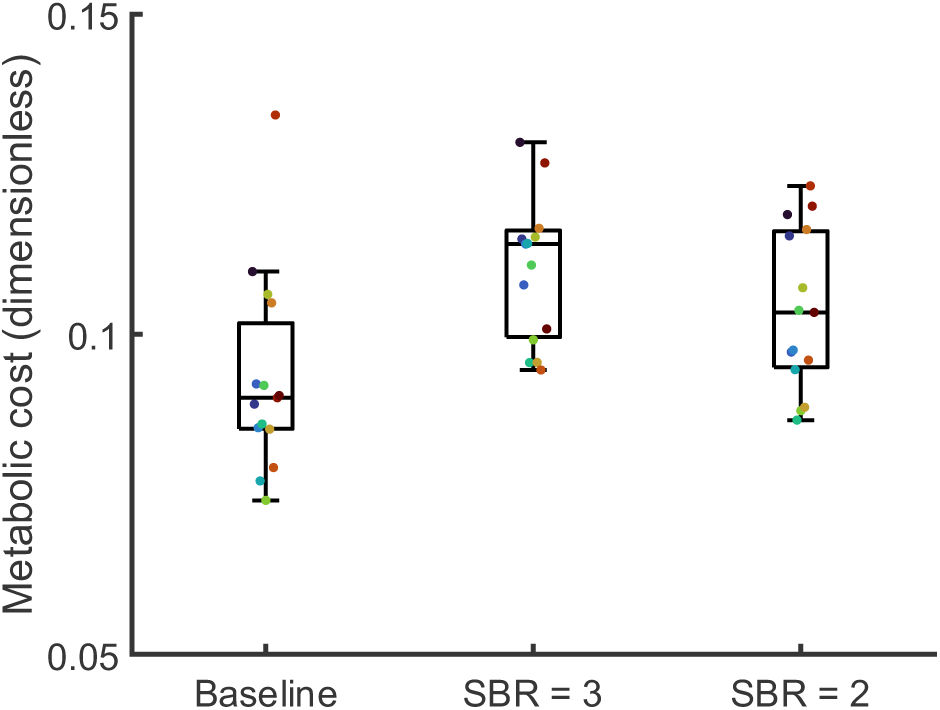
Dimensionless metabolic cost comparison between tied-belt baseline and the free exploration trials with split-belt ratios (SBR) of 3 and 2. Each data point represents a single free exploration trial of a single participant, and different colors represent different participants. Within each box, the center line indicates the median, while the box represents the inter-quartile range.

We compared participants’ spatiotemporal parameters between the two free exploration trials during two distinct phases of the trial: early adaptation (EA) and steady state (SS). We defined early adaptation as strides 34 – 44 (the subsequent 10 strides immediately after the 40-second mark averaged across participants) (please see Methods for this definition) and defined steady state as the last 100 strides of the trial.

We calculated and compared seven outcomes: 1) step length on fast belt, 2) step length on slow belt, 3) foot swing distance on fast belt, 4) foot swing distance on slow belt, 5) time spent on fast belt, 6) time spent on fast belt, 7) SLA. For statistical analysis, we applied paired student’s t-test with a Bonferroni correction. The significance level for the overall test was set at p-value of 0.05, and we report the adjusted p-values below.

First, we analyzed if participants adapted their gait parameters during the first 6-minute free exploration trial at the same split-belt ratio as the guided trials (SBR=3). We found no significant differences in any of the outcomes we analyzed between early adaptation and steady state phases (Table 1, Figure 5).

**Figure 5.**
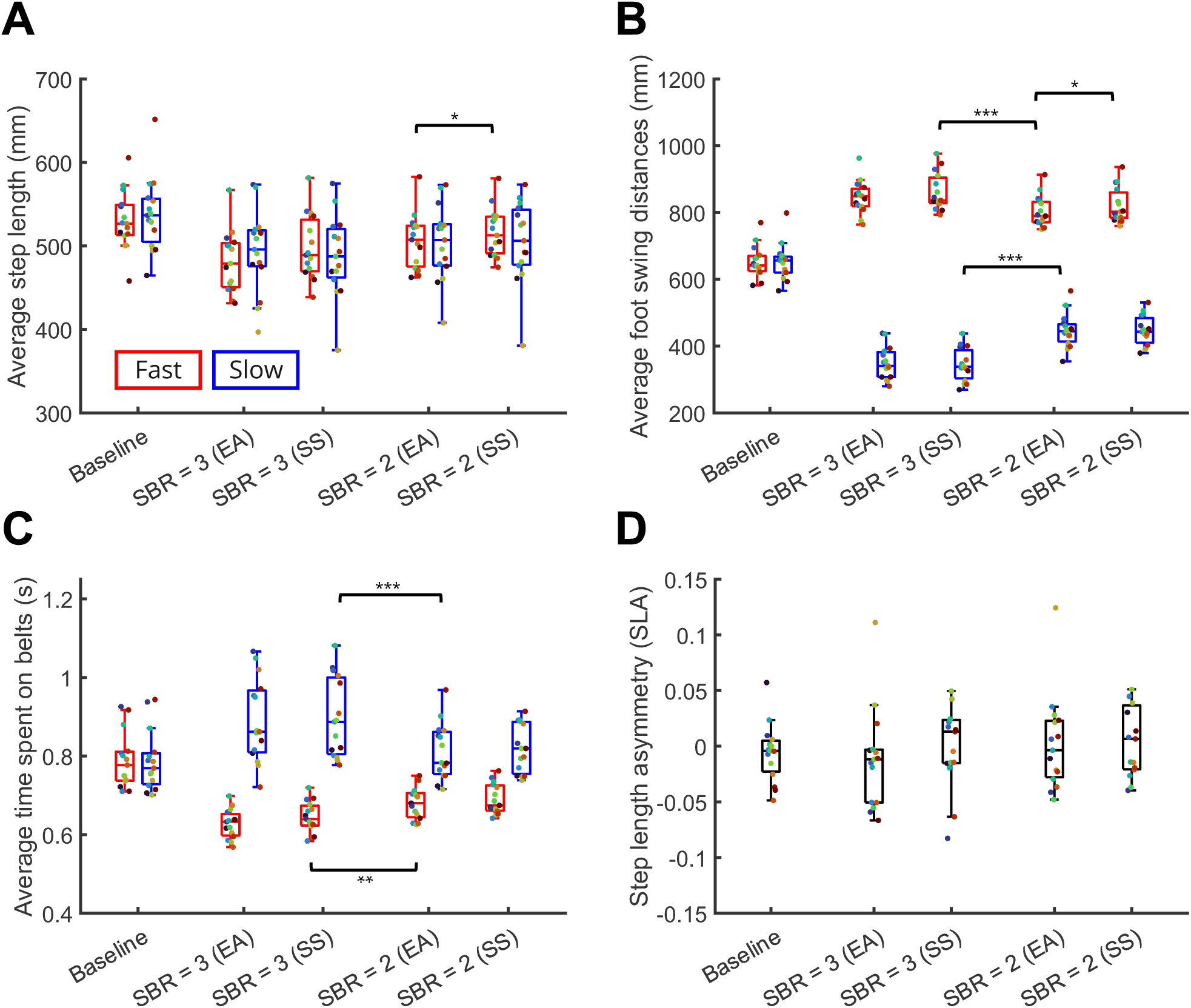
Comparison between tied-belt baseline and the two free exploration trials’ early adaptation (EA) and steady-state (SS) phases for (A) step length, (B) foot swing distance, (C) time spent on each belt, and (D) SLA. For notational consistency, the left and right belts of the tied-belt baseline condition are denoted fast and slow belt, respectively, despite equal belt speeds. Red and blue boxes represent the fast and slow belts, respectively. Within each box, the center line indicates the median, while top and bottom whiskers represent the first and third quartile, respectively. Each data point represents a single free exploration trial of a single participant, and different colors represent different participants. Statistical significance was represented by * (p<0.05), ** (p<0.01) and *** (p<0.001), respectively.

**Table 1.** Summary of biomechanical changes between early exploration (EA) and steady-state (SS) phases of Exploration Trial 1 (SBR = 3) and Exploration Trial 2 (SBR = 2). P-values reported are adjusted per Bonferroni correction for 21 tests conducted.

| Gait Parameter |  | Comparing<br>EA to SS during<br>Exploration Trial 1 | Comparing<br>SS of Exploration Trial 1 to<br>EA of Exploration Trial 2 | Comparing<br>EA to SS during<br>Exploration Trial 2 |
| --- | --- | --- | --- | --- |
| SLA | | No change ($p = 1$) | No change ($p = 1$) | No change ($p = 0.76$) |
| Step<br>length | Fast belt | No change ($p = 0.24$) | No change ($p = 1$) | Increased ($p = 0.04$) |
| | Slow belt | No change ($p = 1$) | No change ($p = 0.17$) | No change ($p = 1$) |
| Swing<br>distance | Fast belt | No change ($p = 1$) | Decreased ($p < 0.001$) | Increased ($p = 0.01$) |
| | Slow belt | No change ($p = 1$) | Increased ($p < 0.001$) | No change ($p = 1$) |
| Time<br>spent on<br>belt | Fast belt | No change ($p = 0.29$) | Increased ($p = 0.005$) | No change ($p = 1$) |
| | Slow belt | No change ($p = 1$) | Decreased ($p < 0.001$) | No change ($p = 1$) |

We then compared participants’ gait parameters between steady state of first free exploration trial (SBR = 3) and early adaptation of the second free exploration trial (SBR = 2) (Table 1). We found no significant difference in SLA between these conditions (p = 1). We found that participants significantly decreased their swing distance on the fast belt (p< 0.001) and time spent on the slow belt (p< 0.001) (Figure 5 B). Conversely, they significantly increased their swing distance on the slow belt (p< 0.001) (Figure 5 B) and time spent on the fast belt (p= 0.005) (Figure 5 C).

During the second free exploration trial with the smaller split-belt ratio (SBR = 2), participants continued to adapt some of their gait parameters over the 6-minute trial, but not all of them. Specifically, we observed no significant changes in time spent on either belt (p = 1 and p = 1, respectively) between early adaptation and steady state phases. Meanwhile, compared to early adaptation, at steady-state participants significantly increased their step lengths on the fast belt (p = 0.04) while maintaining similar step lengths on the slow belt (p = 1). This change was likely due to a significant increase in swing distance on the fast belt (p = 0.01), while swing distance on the slow belt remained unchanged (p = 1). These gait changes together contributed to an increasing (but not statistically significant) trend for SLA between early adaptation (0.002 ± 0.043) and steady state (0.013 ± 0.052) (p = 0.76). (Figure 5 D)

As stated previously, participants did not self-select SLAs that could lead to a metabolic cost reduction despite broad experience (Figure 3). To investigate this observation, we analyzed whether participants were able to reliably achieve the target SLAs prescribed by the real-time visual feedback. To this end, we computed the difference between their achieved steady-state SLA and the target SLA (Figure 6). We found that participants generally under-achieved the prescribed SLAs, walking at less negative SLAs at negative target levels as well as less positive SLAs at positive target levels. Participants were able to most accurately track the target SLAs that were close to symmetry, achieving the smallest absolute errors at SLA targets of −0.05 and 0. Conversely, for the most positive and negative SLA targets, participants performed poorly. Notably, extremely positive SLAs seemed harder to achieve than their negative counterparts, with the most extreme positive SLA target of +0.15 having approximately double the error of the most extreme negative SLA target of −0.15.

**Figure 6.**
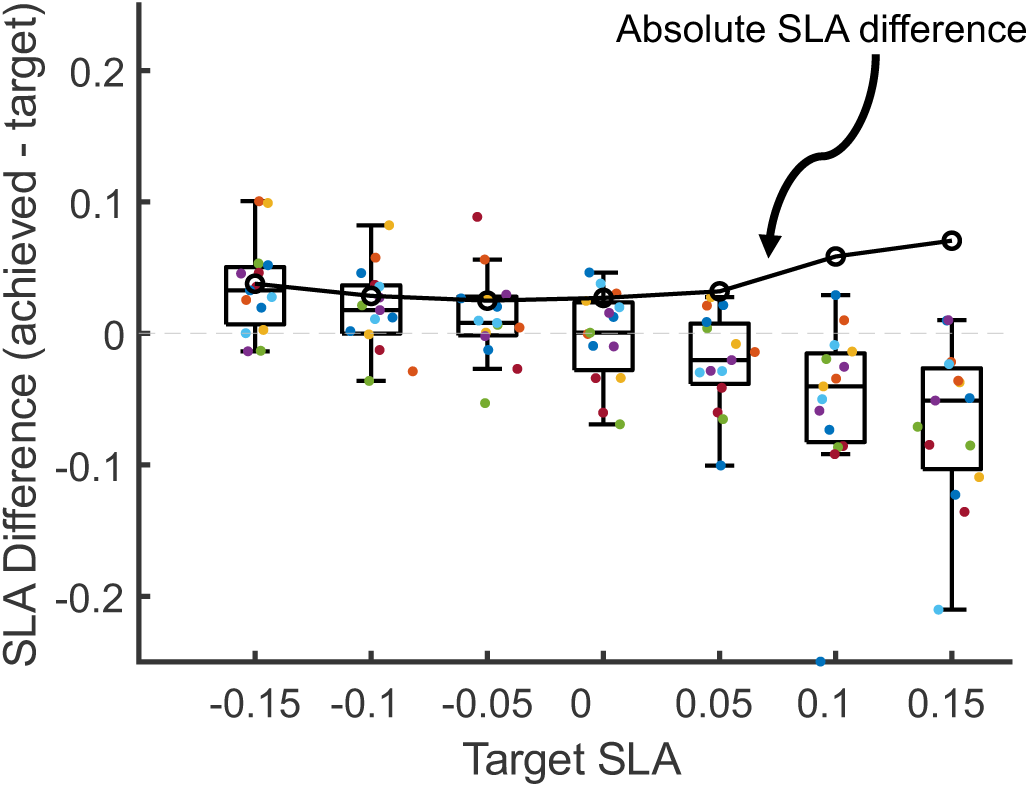
Difference between participants’ achieved SLA and the target SLA during guided exploration trials. Each data point represents a single guided exploration trial of a single participant, and different colors represent different participants. Within each box, the center line indicates the median, while top and bottom whiskers represent the first and third quartile, respectively. The solid black line with circle markers represents the mean **absolute value** of the SLA differences at each target SLA.

## Discussion

The purpose of our study was to analyze how people adapt their gait on a split-belt treadmill when they are asked to walk at a smaller split-belt ratio than the condition in which they gained broad and extensive experience. To this end, we replicated the study protocol from Sanchez et al.^11^ to guide participants to broadly explore the metabolic cost landscape of walking on a split-belt treadmill at a fixed split-belt ratio of 3. Participants were encouraged to walk at different, prescribed step-length asymmetries (SLAs) using real-time visual feedback of their step length. After this guided exploration, participants performed a free exploration trial at the same split-belt ratio, which also follows the protocol established by Sanchez et al.^11^. In our study, we added a second free exploration trial at a smaller split-belt ratio of 2, to quantify how participants might adapt their gait to this new condition that they had not previously experienced.

During the *guided exploration phase* of our experiment, our observations generally matched those reported in Sanchez et al.^11^ We observed that, at positive SLAs, participants’ legs shifted from doing net positive work to doing net negative work, which agrees with theoretical predictions. Notably, participants achieved this by reducing their legs’ positive work rate in response to the increased net positive work rate by treadmill rather than increasing their legs’ negative work rate. This reduction in human positive work at higher SLAs was also correlated with a decrease in metabolic cost (Figure 2D), suggesting that participants were able to accept the positive work done by the treadmill and decrease the positive work done by their legs, ultimately reducing their energy cost. One key difference between our results and those reported in Sanchez et al. is that we observed a *linear* decrease in metabolic cost as a function of SLA, while Sanchez et al. reported a *quadratic* relationship; the second order coefficient in our model was not significantly different from zero (p = 0.29). This indicates that participants in our experiment did not consistently reach an upper bound on the assistance they could receive from the treadmill at higher SLAs. In subsequent analyses, we found that, despite the real-time feedback, our participants were not able to consistently achieve SLA of +0.1 to +0.15 (Figure 6). It is possible that the under-sampling of this higher range of SLAs could explain why we did not observe a second order trend.

Surprisingly, despite the guided exploration protocol, participants did not self-select a more positive SLA than their baseline during either of the free exploration trials. As discussed above, we did observe that metabolic cost and net work done by the human legs significantly decreased with increasing SLA during the guided exploration trials. This means that our participants did in fact receive assistance from the treadmill and experienced gait patterns that incurred lower cost. However, in contrast to Sanchez et al.^11^ and other studies using robotic exoskeletons^28,29^, broad experience within the landscape and exposure to lower cost gaits did not initiate participants’ spontaneous adaptation to energetically favorable asymmetric gait when the external guidance was removed. There are any number of potential explanations for this observed phenomenon. Some have hypothesized that the energetic savings associated with adopting a highly asymmetric gait may be too minor compared to the cost of maintaining symmetry to initiate optimization^33^, but this remains an active area of study in biomechanics and motor learning.

Given this discrepancy, we looked more deeply into the *quality* of the exploration experience. We found a difference between our participants’ ability to follow the prescribed SLAs compared to what was reported in Sanchez et al. As illustrated in Figure 6, our participants were only able to achieve target SLAs near symmetry with precision, while undershooting both highly positive and negative target SLAs. Although participants were shown the real-time position of their feet, they were not instructed exactly how to change their steps to meet the target SLA. Indeed, our participants achieved an average of 6.49 ± 0.41 step score (a perfect score is 10) across all guided trials (see Methods for score computation). As a comparison, Sanchez et al. reported that their participants were “successful at maintaining scores of above 8 points for each stride”^11^. As such, it is possible that our pool of participants had a harder time achieving the prescribed SLAs, which could have increased the perceived effort associated with such SLAs. While we did not measure perceived effort^35^ directly, some participants verbally expressed frustration with being unable to achieve the prescribed SLA. Some prior work has suggested that perceived effort may influence people’s choice of gait parameters^36^, which is one explanation for why our participants did not self-select positive SLAs during the free exploration trials.

To gain insight into whether participants learned a fixed gait pattern or if they continuously adapted their gait in response to a new energetic landscape (i.e., the smaller split-belt ratio), we examined how participants adapted their gait within each of the free exploration trials. Between the early adaptation and steady-state phases of the first free exploration trial (at the same split-belt ratio of 3 as guided exploration), we found no significant changes in any of the biomechanical variables analyzed. This suggests that participants had, throughout the previous guided exploration trials, developed a relatively consistent strategy for walking at this split-belt ratio.

When transitioning from the higher split-belt ratio to a lower one, participants adjusted this strategy. We observed participants significantly altering their foot swing distances as well as time spent on belts for both legs. This is somewhat expected, because maintaining the exact same gait variables from the higher split-belt ratio would have resulted in an extremely asymmetric gait. In this case, participants would likely experience discomfort, difficulty balancing, or trouble remaining on the treadmill due to nonzero net accelerations. Interestingly, the change in foot swing distances and time spent on belts together led to both fast and slow leg step lengths staying relatively unchanged. This is possibly due to participants attempting to maintain their previously developed strategy while staying on the treadmill comfortably.

Throughout the second free exploration trial with the lower split-belt ratio, we observed continued change in some gait variables, but not all of them. Notably, compared to early adaptation period, at steady state participants had increased step lengths on the fast belt, presumably because of increased foot swing distance on this belt. We speculate that these changes suggest that participants actively modified their previously learned strategy, rather than passively adjusting their gait to maintain balance or to stay on the treadmill. Further, we speculate that this strategy modification is at least in part explained by energy optimization, enabled due to broad experience of energy cost landscape at higher split-belt ratio. This idea is partially supported by the fact that participants experienced decreased steady-state metabolic cost when walking at split-belt ratio of 2 compared to walking at split-belt ratio of 3. Therefore, it is possible that participants actively adjusted their gait strategy to adopt a more energetically efficient gait at the lower split-belt ratio.

That said, we cannot rule out the possibility that participants modified their strategy solely because of prolonged exposure to split-belt walking in general. Indeed, Sanchez et al. reported that people spontaneously adapted to a positive SLA while walking on a split-belt treadmill continuously for 45 minutes^18^. However, we believe that this explanation is unlikely because the experimental condition in Sanchez et al. remained constant throughout the prolonged exposure. Meanwhile, multiple studies^21,22^ with non-static conditions, whose total walking time exceeded this time range when summed together over trials, did not report consistent spontaneous adaptation, suggesting that prolonged exposure in non-static conditions might not initiate the same level of adaptation.

Similarly, while we speculate that energy optimization is a potential explanation for why participants modified their strategies, the decreased metabolic cost observed in the second free exploration trial at the smaller split-belt ratio may not serve as definitive evidence. This is because walking at different split-belt ratios creates different energy landscapes. Indeed, Butterfield and Collins have proposed that when average belt speed was held constant, an increased difference in belt speeds can be modeled as an increase in equivalent tied-belt speeds, even though the effect is relatively small^21^. This effect by itself would result in increased metabolic demand as split-belt ratio increases. Therefore, although we did observe lower average metabolic cost for the second, smaller split-belt ratio compared to the first, larger split-belt ratio, we cannot confirm whether the observed gait adaptations led to additional energy optimization. One potential way to further investigate this is to compare the participants’ metabolic cost between early adaptation and steady state at both split-belt ratios. However, due to the delayed nature and stabilization time requirement of indirect calorimetry, we were unable to do so with this protocol.

The main purpose of our study was to analyze how people adapt to a smaller split-belt ratio after experiencing and adapting to a larger one. Therefore, we designed the free exploration trials at split-belt ratio of 2 immediately after free exploration trials at split-belt ratio of 3. Because we were mostly interested in how people adapt their existing strategies as opposed to learning a new one, we did not include tied-belt wash-out trials as other studies evaluating learning or retainment often do^8,24,37^. For the same reason, we did not include a full set of guided exploration trials at a split-belt ratio of 2. Future work could explore if the order of exposure to different split-belt ratios plays a role.

Here, we argued that the significant change in some gait variables during free exploration at split-belt ratio of 2 that weren’t present during free exploration at split-belt ratio of 3 suggests that our participants adapted their gait. However, these observations happened after an apparent lack of spontaneous adaptation to walking in positive SLA in both free exploration trials. It is currently unclear what the implications are of this absence. Without deeper understanding of the mechanisms behind the adaptation itself, we cannot speculate what would happen had we analyzed a cohort who did initiate spontaneous adaptation to positive SLA. Additionally, the positive correlation between split-belt ratio and human metabolic cost we observed suggested that contrary to theoretical predictions, our participants weren’t consistently able to exploit the potential assistance from the split-belt treadmill. These results are consistent with previous *in vivo* results^21,22^, and suggest that human adaptation on split-belt treadmill is a more nuanced process than simply an energy-optimization-based model. Future work could explore the physiological mechanisms governing spontaneous adaptation (or lack thereof) to broad guided experience. Such research could elucidate the contributions of different factors, such as energy optimization and center of mass stabilization.

## Methods

### Participants

We recruited 15 participants (9 male and 6 female) with no prior experience walking on a split-belt treadmill (mean ± SD age: 27.1 ± 4.3 years; height: 174.2 ± 10.5 cm; mass: 76.1 ± 14.5 kg; leg length 83.3 5.31 cm) for this study. Participants self-reported their gender, height and mass, and we measured leg length as the distance from the lateral malleolus to trochanter. All participants reported no current or past cardiovascular, pulmonary, neurological, or musculoskeletal conditions which prevent them from performing mild-to-moderate exercise. All study procedures were approved by the University of Washington Internal Review Board (STUDY00020166). All participants provided written, informed consent.

### Data Acquisition

Participants walked on a split-belt instrumented treadmill (Bertec Corporation, Columbus, OH, USA). Ground reaction force data were collected at 1000 Hz with treadmill-integrated force plates. Participants’ ankle positions were measured using retro-reflective markers placed on the lateral malleoli of each foot, tracked by a 12-camera Qualisys Oqus Motion Capture system (Qualisys AB, Goteborg, Sweden) at 100 Hz. Metabolic cost was measured using indirect calorimetry. Rates of oxygen and carbon dioxide consumption were measured breath-by-breath using the COSMED K5 wearable metabolic system (COSMED, Rome, Italy).

### Study Design

All participants fasted for at least 3 hours before the start of the experiment for consistency in metabolic cost measurement. A graphical experimental protocol is shown in Figure 1A. We first measured participants’ baseline metabolic cost during a 6-minute period of quiet standing. We began the walking trials with a tied-belt trial, with both belts of treadmill running at 1 m/s (*baseline*), during which we recorded the participants’ baseline stride length and step length asymmetry (SLA). Participants then performed another tied-belt trial with real-time feedback (*baseline with feedback*), with target SLA equal to their baseline SLA. Details about the visual feedback are provided in the next section.

Subsequently, participants performed seven *guided exploration* trials: split-belt trials with real-time feedback and a split-belt ratio of 3 (fast and slow belt speeds were 1.5 m/s and 0.5 m/s, respectively). Participants were encouraged to walk at different SLAs using real-time visual feedback. The prescribed SLAs included those which might penalize or benefit the metabolic cost of walking (−0.15, −0.1, −0.05,0, +0.05, +0.1, +0.15). We randomized the order of trials for each participant. All guided exploration trials lasted 6 minutes each.

Following the *guided exploration* trials, participants performed a *free exploration* trial at the same split-belt ratio 3, followed by another *free exploration* trial at a smaller split-belt ratio of 2 (fast and slow belt speeds were 1.33 m/s and 0.67 m/s respectively). All free exploration trials were conducted without real-time feedback and lasted 6 minutes each. We verbally encouraged participants to walk freely and comfortably.

Participants rested for at least 4 minutes between walking trials. We ensured that their metabolic rate had returned to resting level before starting a new trial by visual inspection. The left belt was the fast belt and the right belt was the slow belt across all split belt trials.

### Real-Time Feedback

We replicated the experimental protocol from Sanchez et al.^11^ to guide participants to broadly explore the energy landscape associated with different SLAs. During *baseline with feedback* and all *guided exploration* trials, an iPad positioned at the participant’s eye level provided them with real-time information about their foot positions. The monitor displayed one bar for each leg, with the bar height corresponding to the target step lengths needed to achieve the prescribed SLA (Figure 7).

**Figure 7.**
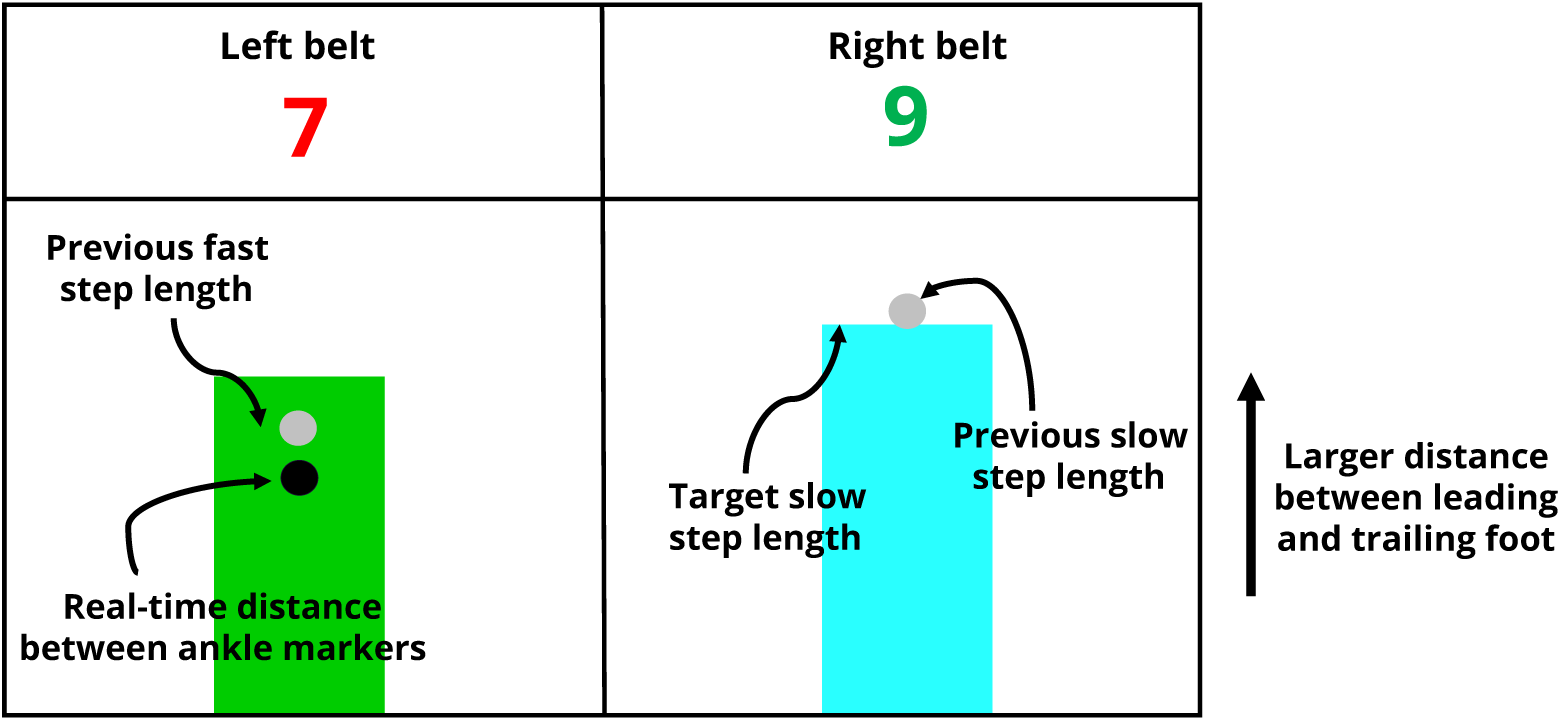
The GUI displayed visual feedback to the participant about the real-time distance between their feet (black dot), the target step length for each foot (green and cyan bars), and the length of their previous step (gray dot). It also displayed a score indicating how well they were matching the target; scores greater than 8 were green, and less than 8 were red.

We computed the target step lengths as follows: for all trials with feedback, the sum of fast and slow leg target step lengths were held constant and equal to the sum of each subject’s baseline step lengths. During the *guided exploration* trials, we back computed the target step lengths for fast leg and slow leg to achieve the desired SLA. During the *baseline with feedback* trial, both left and right target step lengths were set to the participants’ baseline step length.

We defined heel strikes for each leg when the vertical ground reaction force on the corresponding belt surpassing a threshold of 30 N. We defined strides from the heel strike on the fast belt to the subsequent heel strike on the fast belt. We calculated step lengths (𝑆𝐿) for each leg as the instantaneous distance between the participant’s ankle markers at heel strike. 𝑆𝐿_𝑓𝑎𝑠𝑡/𝑠𝑙𝑜𝑤_ was defined when the fast/slow leg was the leading leg at heel strike. We computed SLA for each stride according to the following equation^10,11,22^:

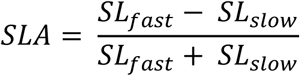

The instantaneous anterior-posterior distance between the participant’s leading and trailing ankle marker was computed and displayed by a black dot moving along the bar corresponding to the leading leg. This distance was used to calculate step lengths at heel strike, so participants were shown in real-time how large their step length would be if they were to take a heel strike. Once heel strike occurred, the black dot disappeared, leaving a gray mark indicating the actual step length achieved. A score was also displayed on top of the bar at each heel strike to better inform the participants of their performance. This score was computed according to the following equation^11^:

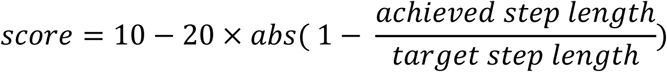

The score was displayed in green if the score was 8 or higher, red if the score was between 2 and 8, and disappeared otherwise. We verbally instructed participants to try to maintain a green score and encouraged them to aim at a score of 10 at every step.

### Post-hoc Analysis

For post-hoc analysis, ground reaction forces and marker position data were low-pass filtered with a fourth-order Butterworth filter with cutoff frequency of 20 Hz and 10 Hz, respectively. In addition to step lengths and SLA, in post-hoc analysis we defined toe-offs for each leg as vertical ground reaction force on the corresponding belt decreasing below the same threshold of 30 N. We computed *time spent on belt* for each leg as the time between heel strike and toe-off for the corresponding leg, and *foot swing distance* as the anterior-posterior distance between toe-off and subsequent heel strike.

Any segmented strides that did not include exactly one fast step and one slow step were discarded from the analysis. We averaged the last 100 strides of each trial for computing steady-state SLA and step lengths. We defined the first 10 strides immediately following the 40 seconds mark during free explorations as “early adaptation” (EA). Previous studies have shown that gait adaptation can be modelled to occur at two distinctive time scales^38^, with the fast time scale associated more with stability and slow time scale associated more with energy optimization^19,20^. A recent publication found that during split-belt walking, the margin of stability converged quickly for participants at a single fast time scale, with a time constant of around 40 seconds^25^. We therefore defined the 10 strides that participants on average walked immediately after the 40-second mark as the early adaptation phase. This translated to steps 34 – 44 for each exploration trial, which we averaged to compute early adaptation SLA and other biomechanical variables.

While the real-time feedback encouraged participants to walk at preset, discrete SLAs, we used the real SLAs achieved by participants, which was a continuous variable, for biomechanical analyses.

We computed work rate performed by participants’ legs and the treadmill using a modified individual limbs method that had previously been applied to analyze energetics during split-belt walking^11,12,16,34^. Briefly, this method assumes legs to be massless pistons that apply equal but opposite direction forces to the treadmill and human’s center of mass (COM), simplified to be a point mass. We performed the analysis on a stride-by-stride basis. Within each stride cycle, the velocities of COM in each direction were computed by integrating the accelerations due to forces acting on COM. We then computed work rates from the dot product of velocities and forces. We summed positive work rates and negative work rates throughout a stride cycle to compute the net work rate.

For metabolic cost, we assumed the last three minutes’ data of each trial to be at steady-state^28^.

We then computed steady-state metabolic cost computed using the Brockway equation^39^ as following:

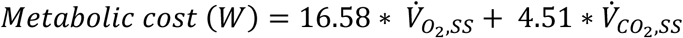

Where 𝑉̇_𝑂2,𝑆𝑆_ and 𝑉̇_𝐶𝑂2,𝑆𝑆_ are the average steady-state volumetric consumption and generation rate (mL/s) of oxygen and carbon dioxide, respectively. Net metabolic cost was computed by extracting subject-specific standing metabolic cost from each trial’s values.

Lastly, to minimize inter-participant variability, we normalized metabolic cost and mechanical work rates to dimensionless units (as previously described^11^) by dividing each individual’s values by 𝑚𝑙^0.5^𝑔^1.5^, where 𝑚 is the body mass (kg), 𝑙 the leg length (m) and 𝑔 the gravitational constant (9.8 𝑚/𝑠^2^).

## Statistical Analysis

To account for individual factors affecting their energetic and metabolic cost, we built linear mixed effect models (LMEM) using Matlab *fitlme* function to analyze work and work rates by human legs and the treadmill as well as metabolic cost ^11^. Specifically, we built LMEMs to capture the relationships between: 1) net work rate done by human legs and human SLA, 2) net work rate done by the treadmill on human legs and SLA, 3) positive work rate done by human legs and net work rate done by the treadmill on human legs, and 4) human metabolic cost as a function of SLA. All these models included the independent variable as a fixed effect and a random effect intercept for each participant. For visualization purposes only, we removed the random intercept from each data point in the figures.

To compare participants’ biomechanical variables between free exploration trials at split-belt ratio of 3 and 2, during early adaptation and steady-state, respectively, we applied paired student’s t-test with Bonferroni correction to account for multiple comparisons. Specifically, we compared the following variables: 1) step length on fast belt, 2) step length on slow belt, 3) foot swing distance on fast belt, 4) foot swing distance on slow belt, 5) time spent on fast belt, 6) time spent on fast belt, and 7) SLA. Each variable was compared across three conditions: i) between EA and SS of SBR = 3, ii) between SS of SBR = 3 and EA of SBR = 2, and iii) between EA and SS of SBR = 2, for a total of 7x3 = 21 comparisons. Overall significance level was set at p-value of 0.05, and all the reported p-values for this analysis were adjusted by multiplying 21.

All other significance levels were set at p-value of 0.05.

All statistical analyses were performed in Matlab 2023b (MathWorks, Natick, MA).

## Data Availability

All data collected during this experiment have been processed as described in the manuscript and are freely available via Dryad (DOI: 10.5061/dryad.rbnzs7hr4). The code used to analyze and generate the results in this manuscript using these data is provided in GitHub at https://github.com/ingraham-research/Split-belt-paper-2025.

## Acknowledgements

We thank Annika Pfister for comments and feedback. We thank all the participants who completed this research study for their time.

## Author Contributions

Z.J. contributed to the design of experiments, performed experiments, collected and analyzed the data. J.I and S.A.B. contributed to the design of experiments. K.A.I. contributed to the design of experiments and analysis and interpretation of data. All authors contributed to drafting the manuscript. All authors read and approved of the final manuscript.

